# Symbiont community diversity is more constrained in holobionts that tolerate diverse stressors

**DOI:** 10.1101/572479

**Authors:** Lauren I. Howe-Kerr, Benedicte Bachelot, Rachel M. Wright, Carly D. Kenkel, Line K. Bay, Adrienne M.S. Correa

**Author notes:** **Corresponding author:** Lauren I. Howe-Kerr, 6100 Main Street, MS-140, Houston, TX 77005, ORCID: 0000-0002-8086-5869.

## Abstract

Coral reefs are experiencing global declines as climate change and other stressors cause environmental conditions to exceed the physiological tolerances of host organisms and their microbial symbionts (collectively termed the holobiont). To assess the role of symbiont community composition in holobiont stress tolerance, diversity metrics and abundances of obligate dinoflagellate endosymbionts (Family: Symbiodiniaceae) were quantified from eight *Acropora millepora* coral colonies (hereafter called genets) that thrived under or responded poorly to various stressors. Four ‘best performer’ coral genets were selected for analysis because they survived 10 days of high temperature, high *p*CO_2_, bacterial addition, or combined stressors, whereas four ‘worst performer’ coral genets were analyzed because they experienced significant mortality under these stressors. At the end of the experimental period, seven of eight coral genets mainly hosted *Cladocopium* symbionts, but also contained *Brevolium, Durusdinium*, and/or *Gerakladinium* symbionts at lower abundances (<0.1% of the total community). After 10 days of stress, symbiont communities varied significantly among host genets, but not stress treatments, based on alpha and beta diversity metrics. A generalized joint attribute model (GJAM) also predicted that symbiont communities were primarily sensitive to host genet at regional scales. Indicator species analysis and the regional GJAM model identified significant associations among particular symbionts and host genet performance. Specifically, *Cladocopium* 3k contributed to the success of best performer host genets under various stressful conditions, whereas *Durusdinium glynnii* and *Durusdinium trenchii* were significantly associated with one worst performer genet. *Cladocopium* 3k dominance should be more broadly investigated as a potential predictor of stress resistance in *Acropora millepora* populations across their geographic range. Symbiodiniaceae communities exhibited higher richness and variance (beta diversity) in the worst performing genets. These findings highlight that symbiont community diversity metrics may be important indicators of resilience in hosts central to diverse disciplines, from agriculture to medicine.

## Introduction

Coral reefs are undergoing rapid declines in health on a global scale following increased exposure to climate change stressors, such as warming sea surface temperatures and *p*CO_2_, and increased loading with disease-causing agents (Hughes et al., 2017, 2018; Zaneveld et al., 2016). Microbial symbionts can influence the capacity of hosts, including reef-building corals, to acclimatize to environmental stressors. Dinoflagellates in the family Symbiodiniaceae reside in the tissues of corals, giant clams, and other marine invertebrates, and have been empirically demonstrated to influence the ability of corals to survive stress events (Baker, 2001; Baker, Starger, McClanahan, & Glynn, 2004; Kenkel & Bay, 2018; Todd C. LaJeunesse et al., 2010; Todd C. LaJeunesse, Smith, Finney, & Oxenford, 2009; Manzello et al., 2018; Rouzé, Lecellier, Saulnier, & Berteaux-Lecellier, 2016). Particular Symbiodiniaceae species, such as *Durusdinium trenchii*, are more likely to remain associated with hosts and/or to retain photosynthetic function during acute temperature anomalies or extremes (Manzello et al., 2018; Silverstein, Cunning, & Baker, 2015; but see Silverstein et al., 2017). For example, in a dominant reef-building Pacific coral *(Acropora millepora)*, a *Durusdinium* species minimized bleaching (a diminished host health state characterized by loss of Symbiodiniaceae *en masse)* and increased host survival of acute thermal anomalies (Bay, Doyle, Logan, & Berkelmans, 2016; Berkelmans & van Oppen, 2006; A. M. Jones, Berkelmans, van Oppen, Mieog, & Sinclair, 2008; Mieog et al., 2009). Yet, harboring these stress resistant symbionts might constitute a tradeoff for corals: increased temperature tolerance at the expense of growth especially in cooler environments (Cantin, van Oppen, Willis, Mieog, & Negri, 2009; A. Jones & Berkelmans, 2010; Little, van Oppen, & Willis, 2004). Elevated *p*CO_2_ has been shown to enhance growth and photosynthetic capacity in certain Symbiodiniaceae species (Brading et al., 2011), to have no effect on others (Brading et al., 2011) or to result in the loss of symbionts and their photosynthetic function (Kaniewska et al., 2012). Thus, the impact of ocean acidification stress on Symbiodiniaceae likely varies with taxa and context and needs further investigation. Interactions between Symbiodiniaceae and bacterial pathogens (e.g., *Vibrio* spp.) have also previously been correlated with coral (Rouzé et al., 2016) or symbiont health (Hauff et al., 2014). Specifically, *Acropora cytherea* colonies harboring *Durusdinium* symbionts were more resistant to infection with *Vibrio* spp. than conspecifics harboring *Symbiodinium* symbionts (Rouzé et al., 2016). However, the relationship between Symbiodiniaceae identity and relative resistance to *Vibrio* infection is unexplored for most Symbiodiniaceae species, as is the relationship between symbiont community diversity metrics and host resistance to bacterial pathogens.

To date, most Symbiodiniaceae studies, including many of those discussed above, have focused on the contributions of individual symbiont species, rather than symbiont community composition, to holobiont stress responses. This focus invokes the ‘selection effect’, which assumes that the independent function of dominant species drives overall community function (Loreau et al., 2001). However, if the function of a given symbiont species differs when in the context of a diverse community (due to facilitative or other interactions – the ‘complementarity effect’, Fox, 2005), as compared to in monoculture, then community composition or diversity metrics may better represent symbiont contributions to holobiont physiology. For example, community diversity metrics may provide important insights in systems where low abundance symbionts contribute (even if ephemerally) to overall holobiont stress responses (e.g., LaJeunesse et al., 2009; Zaneveld, McMinds, & Thurber, 2017, but see Lee et al., 2016). One of the first studies to investigate how symbiont attributes are associated with host performance (Kenkel & Bay, 2018) found a decreasing trend in symbiont cooperation (in terms of autotrophically derived carbon shared with hosts) under heat stress within coral species that harbored more diverse Symbiodiniaceae communities. Another study found that coral juveniles that harbored more diverse symbiont communities exhibited lower survival than juveniles with less diverse communities (Quigley, Willis, & Bay, 2016). There has also been a general observation that dysbiotic host individuals vary more in microbial community composition than healthy host individuals – the ‘Anna Karenina Principle’ or AKP (Zaneveld et al., 2017). This has previously been documented in experimentally stressed coral-associated bacterial communities (Zaneveld et al., 2016), but has not been examined in the context of Symbiodiniaceae communities.

By identifying symbiont species occurring at abundances as low as 0.1% in a host (Quigley et al., 2014), high-throughput sequencing (HTS) methods make it possible to compare symbiont community characteristics associated with host genets under ambient and stressful conditions, and can support future investigations into the relevance of the complementarity effect for host-microbe symbioses. A variety of markers, including the highly variable Symbiodiniaceae Internal Transcribed Spacer-2 (ITS-2) region of rDNA, have been applied in isolation or combination to assess the diversity of Symbiodiniaceae present in hosts. HTS of the Symbiodinianceae ITS-2 is useful for investigating symbiont contributions to host health and stress response at a fine scale (i.e., at the level of sequence variants, including those present at low abundances); this approach has revealed novel Symbiodiniaceae variants, host associations, and/or distribution patterns (Brener-Raffalli et al., 2018; Cunning, Gates, & Edmunds, 2017; Green, Davies, Matz, & Medina, 2014; Hollie M Putnam, Stat, Pochon, & Gates, 2012; Quigley, Bay, & Willis, 2017; Quigley et al., 2014, 2016; Quigley, Willis, & Bay, 2017; Ziegler et al., 2017; Ziegler, Eguíluz, Duarte, & Voolstra, 2018; Ziegler, Stone, Colman, Takacs-Vesbach, & Shepherd, 2018). For example, variation in the abundances of background (<10% of total community) symbionts has been documented across reefs separated by as little as 19 km (e.g., Green et al., 2014; van Oppen et al., 2018). It has been hypothesized that these differences in Symbiodiniaceae sequence variant distribution may be associated with fine scale environmental variation (Brener-Raffalli et al., 2018; Cunning et al., 2017; Quigley et al., 2014; Quigley, Warner, Bay, & Willis, 2018). However, no HTS studies have examined symbiont community composition or diversity metrics (i.e., alpha and beta diversity) from host genets that perform well versus poorly under stressful conditions.

Here, we applied a HTS approach to determine whether particular Symbiodiniaceae variants (potentially species) or community characteristics were associated with host genets and/or environmental stressors. At the conclusion of experimental stress treatments, we analyzed Symbiodiniaceae communities from eight *Acropora millepora* genets that thrived under or responded poorly to climate stressors (high temperature and/or *p*CO_2_), increased exposure to a pathogenic bacteria *(Vibrio owensii)* in isolation, or when these stressors were combined. Coral fragment health, Symbiodiniaceae cell density, and Symbiodiniaceae identity and diversity metrics were assessed in order to identify symbiont ‘species’ and community diversity attributes that were associated with host survival or mortality. We hypothesized that: 1) best performing host genets will contain higher relative abundances of stress-tolerant Symbiodiniaceae (e.g., *Durusinium trenchii*) than worst performing genets; 2) fragments of a given host genet exposed to different stress treatments will differ significantly in their symbiont community composition and diversity, as well as from that of control fragments; and 3) worst performing host genets will have higher beta diversity than best performing ones, as predicted by the Anna Karenina Principle.

## Materials and Methods

### Study site, experimental design, and sample collection

Colonies representing 8 distinct genets of *Acropora millepora* known to be best or worst performers under stress (based on survival, Wright et al. *submitted)* were collected between October 1-8, 2014 from three Great Barrier Reef sites (Supp. Mat. Figure 1): Pandora Island (18°48’45’’S, 146°25’59.16″, Genets Worst-27, Worst-31, Worst-34), Rib Reef (18°28’53.4’’S, 146°52’24.96″, Genets Best-12, Best-20, Best-38), and Davies Reef lagoon (18°30’3.96’S, 147°22’48″, Genets Best-4, Worst-2) and transferred to flow-through seawater tanks at the National Sea Simulator system at the Australian Institute of Marine Science. Each colony was then split into fragments, acclimatized to a common garden condition for five months, and placed into treatment conditions simulating environmental stressors (Table 1, see the Supplementary Materials for details of the experimental methods).

**Table 1.** Overview of experimental treatments. Eight genets of *Acropora millepora* (N = 120 fragments total, with 3 replicates per genet per treatment), were exposed to control, bacteria addition, elevated temperature, elevated *p*CO_2_, or combined bacteria/heat/*p*CO_2_ stressors. After 10 days, surviving fragments were sampled for Symbiodiniaceae density and diversity.

| Stress type | Temperature<br>(°C) | $p\text{CO}_2$<br>(ppm/pH) | +/- *Vibrio owensii*<br>6 hrs at $10^6$ cells/ml |
| --- | --- | --- | --- |
| Control | 27 | 400 / 8.0 | - |
| Bacteria | 27 | 400 / 8.0 | + |
| Heat | 30 | 400 / 8.0 | - |
| $p\text{CO}_2$ | 27 | 700 / 7.8 | - |
| Combined | 30 | 700 / 7.8 | + |

Here, we analyze the Symbiodiniaceae communities from surviving fragments (N = 102) of four best performing (Best-4, Best-12, Best-20, Best-38) and four worst performing coral genets (Worst-2, Worst-27, Worst-31, Worst-34) following 10 days of exposure to bacteria addition, elevated temperature, elevated *p*CO_2_, or combined bacteria/heat/*p*CO_2_ stressors (samples are a subset of those from Wright et al. *submitted).* Coral fragments were visually inspected and photographed with a Color Watch Card (www.coralwatch.org) every 24 hours for lesions and bleaching signs. Change in fragment color (a proxy for symbiont density and/or chlorophyll content) and buoyant weight growth rate (% change in weight g · day^-1^) were calculated from the beginning to the end of the experiment (see Supplementary Materials for details). At the end of the experiment, surviving coral fragments were snap frozen in liquid nitrogen and then stored at −80°C until processing.

Snap frozen coral fragments were airbrushed to remove coral holobiont tissue, and aliquots of the resulting tissue slurry were transferred to 70% ethanol for DNA extraction or fixed with 5% formalin for Symbiodiniaceae cell density counts. Cell density samples were counted using a haemocytometer (N=8 replicate counts/sample), and density was calculated as cells/cm^2^ by using surface area of the coral skeleton (Stimson & Kinzie, 1991) and total volume of coral tissue blastate. Statistical differences for cell density, color change, and growth rates between treatments and genets were determined using Kruskal-Wallis and Pairwise Wilcoxon rank sum tests.

### Sample DNA extraction and sequencing

DNA from each fragment was extracted using Wayne’s Method (Lundgren, Vera, Peplow, Manel, & van Oppen, 2013). To extract coral holobiont DNA, each sample was centrifuged and the ethanol was removed. The pellet was transferred to a SDS buffer and then precipitated with KOAc, followed by a series of ethanol washes. Pellets were resuspended in 30μl of 1x TAE and stored at −20°C. DNA concentrations were determined through Quant-iT PicoGreen dsDNA assays; 96 samples had sufficient DNA concentrations for HTS sequencing. The ITS-2 region of Symbiodiniaceae rDNA was amplified using symbiont-specific primers: SYM_VAR_5.8SII (5’GAATTGCAGAACTCCGTGAACC 3’) and SYM_VAR_REV 5’ (CGGGTTCWCTTGTYTGACTTCATGC 3’). The target amplicon was approximately 234-266bp (Hume et al., 2018). The PCR reaction contained 5μl of DNA (5ng/μl), 2.5μl of SYM_VAR_5.8SII + MiSeq Adapter (5’-TCGTCGGCAGCGTCAGATGTGTATAAGAGACAG GAATTGCAGAACTCCGTGAACC3’) (2 uM), 2.5μl of SYM_VAR_REV + MiSeq Adapter (5’ -GTCTCGTGGGCTCGGAGATGTGTATAAGAGACAG CGGGTTCWCTTGTYTGACTTCATGC 3’) (2 uM), 12.5μl 2x KAPA HiFi HotStart ReadyMix, and 2.5μl of molecular grade water for a total reaction volume of 25μl. PCR cycles were as follows: 95°C for 3 min, 15 cycles of 95°C for 30 sec, 56°C for 30 sec, and 72°C for 30 sec, and lastly 72°C for 4 min.

PCR clean-up was completed using Agencourt AMPure XP Magnetic Beads. Illumina indexing primers were added to 50μl of purified PCR product, and a new PCR was run to incorporate unique barcodes for each sample. The PCR reaction contained 5μl of cleaned PCR product, 5 μl of Illumina Indexed Primer 1 (i5), 5 μl of Illumina Indexed Primer 2 (i7), 25 μl 2x KAPA HiFi HotStart, and 10 μl PCR Grade water for a total reaction volume of 50μl. PCR cycles were as follows: 95°C for 3 min, 20 cycles of 95°C for 30 sec, 56°C for 30 sec, and 72°C for 30 sec, and lastly 72°C for 4 min. The resulting PCR product was purified with Agencourt AMPure XP Magnetic Beads. Samples were quantified via qPCR using the KAPA library quantification kit and normalized and pooled in equal molar amounts. Pooled samples were sequenced on the Illumina MiSeq platform using a PE300 run with 25% PhiX at the Georgia Genomics and Bioinformatics Core (University of Georgia, Athens, GA).

### Bioinformatic Processing and Statistical Analyses

Illumina’s real time analysis was run during sequencing using the default settings in order to remove clusters with the least reliable data. Demultiplexed fastq files were generated with Illumina’s BaseSpaceFS (version 1.5.964) and reads were processed in RStudio (version 1.1.456) through the DADA2 pipeline (version 1.11.0, Callahan et al., 2016) with modifications for the Symbiodiniaceae ITS-2 region. Samples with less than 10,000 reads (N = 2) were removed from the dataset (N = 94 samples remained). The DADA2 pipeline generated a table of amplicon sequence variants (ASVs); alpha diversity was calculated using Shannon’s Diversity (H’) index and Simpson’s Diversity (1-D) index from the full ASV table. Since these data were not normally distributed based on a Shapiro-Wilk test (H’: W = 0.97, p = 0.01; 1-D: W = 0.89, p<0.001), Wilcoxon rank sum tests were used to test for significant differences in Symbiodiniaceae diversity between best and worst performers, and among treatments. A Kruskal-Wallis test was used to determine whether significant differences existed in the alpha diversity of Symbiodiniaceae ITS-2 sequence variants among treatments.

The ASV table from DADA2 was then curated via the LULU pipeline, which uses co-occurrence patterns and sequence similarity to eliminate erroneous ASVs (note that authentic singleton ASVs could also be removed, which is why alpha diversity statistics were performed prior to the LULU pipeline, Frøslev et al., 2017). Symbiodiniaceae species or ITS-2 types (hereafter, collectively referred to as species) were then assigned based on BLAST results to a local Symbiodiniaceae ITS-2 database (Cunning et al., 2017). Permutational multivariate analysis of variance using Bray-Curtis distance was used to generate an NMDS plot to visualize differences in Symbiodiniaceae community. To determine if Symbiodiniaceae community composition significantly differed between control versus treatment conditions (as a binary comparison), among treatment conditions, between best and worst performing genets, and among treatments within best and worst performing genets, we first tested for homogeneity in group (genets and treatments) dispersion using ‘betadisper’ in the ‘vegan’ R package (version 2.5.3, Oksanen et al., 2013). A significant result indicates that the groups differ significantly in beta diversity. Multivariate analyses of the variance in composition are sensitive to heterogeneity, but permutational multivariate analysis of the variance (‘adonis’ in ‘vegan’) tends to be less sensitive than other tests (Anderson & Walsh, 2013). Therefore, to test for significant differences in Symbiodiniaceae composition among treatments and genets, we used the ‘adonis’ function in ‘vegan’. Symbiodiniaceae species that were strongly indicative of treatment condition or genet designation were examined using the Indicspecies R package (version 1.7.6, De Cáceres & Legendre, 2009). To investigate the strength of links among *A. millepora* genet, stress type, and Symbiodiniaceae community composition at a regional (among reefs) scale, we fitted a generalized joint attribute model (GJAM) in R to the LULU-curated dataset, using 10,000 iterations and 500 burn-in steps (Clark, Nemergut, Seyednasrollah, Turner, & Zhang, 2017). This joint probabilistic model takes into account co-dependence amongst ASVs and performs well in the presence of zeros. We also used the regional GJAM model to inversely predict *A. millepora* genet and experimental treatment given a particular Symbiodiniaceae community. Specifically, for each Symbiodiniaceae sample (*y* *), we used a Monte Carlo integration to inversely predict the covariates (*x* *):

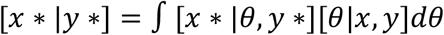

where *θ* is the non-informative prior distribution.

We also estimated the Symbiodiniaceae community sensitivity to host genets and treatment. After fitting the regional GJAM model, we obtained a predictors (genet and treatment)-by-species matrix of coefficients ***B*** that holds all predictor-by-species responses. Elements of ***B*** are the individual sensitivities of each species to each predictor. As part of the fitting process, we also estimated a species-by-species covariance matrix *Σ* that holds residual indirect relationships between species. Sensitivity across the entire Symbiodiniaceae community is then:

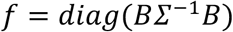

Finally, since Rib Reef genets were all best performers and Pandora Island genets were all worst performers, additional GJAM models were fitted to genets originating from these locations, respectively, to test for treatment effects on more local scales. These local models were identical to the regional GJAM model except that they were fitted only with the data from Pandora Island or Rib Reef.

## Results

### Response to experimental treatments

In response to 10 days of exposure to single or combined stressors, the most coral mortality was observed in the *p*CO_2_ and combined stressors treatments (N = 5 fragments total per treatment, represented as dark grey bars in Figure 1); none of the control fragments experienced mortality.

**Figure 1.**
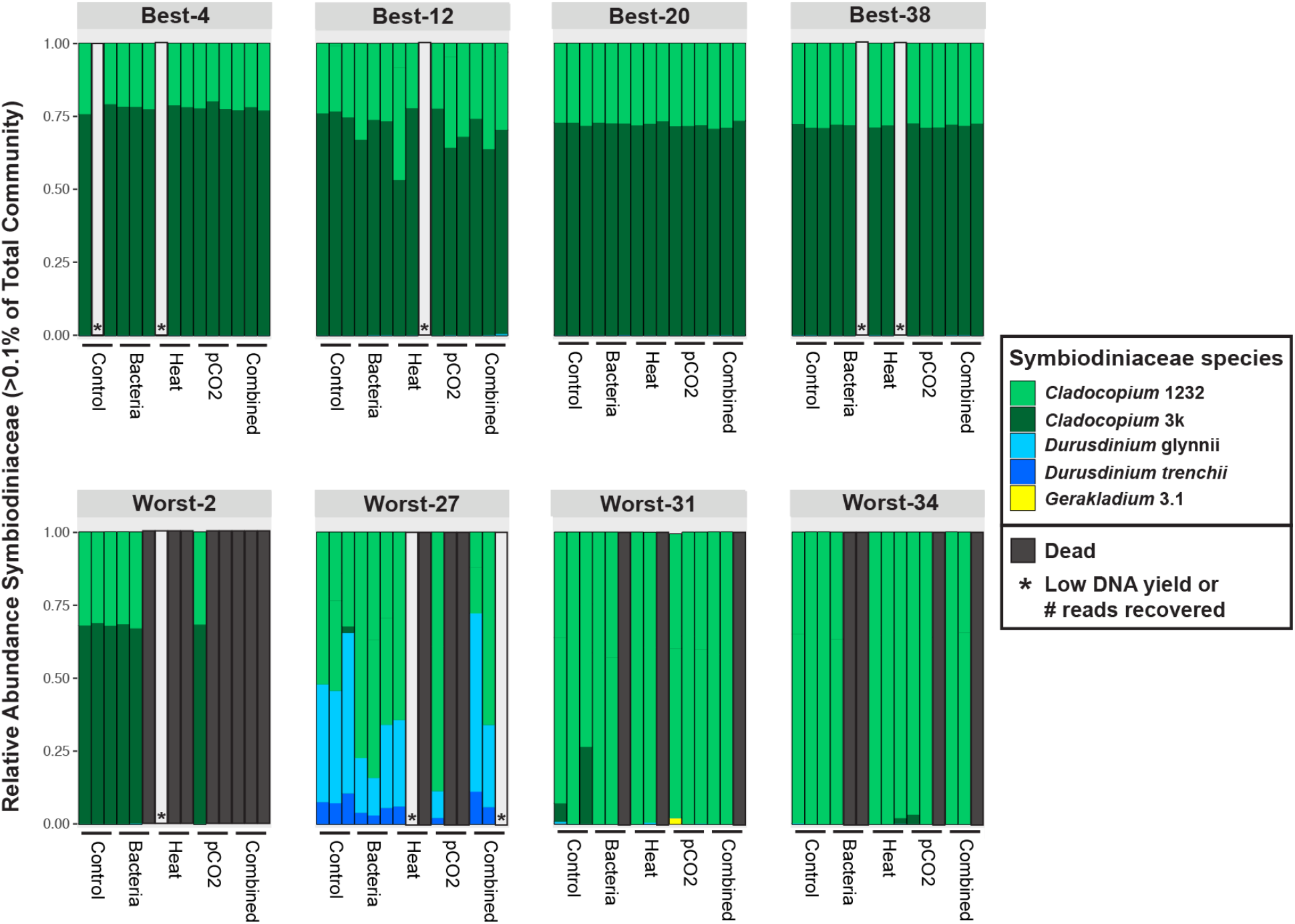
Relative abundance (comprising >0.1% of total community) of Symbiodiniaceae species or Internal Transcribed Spacer-2 (ITS-2) types in individual fragments (vertical bars) of *Acropora millepora* genets exposed to various experimental stressors.

Symbiodiniaceae cell density counts at the end of the experiment ranged from 1.8e^6^ to 1.6e^4^ cells/cm^2^ (average for best performers: 3.5e^5^ + 2.1e^5^cells/cm^2^; average for all worst performers: 4.1e^5^ + 4.0e^5^ cells/cm^2^; Figure 2). Worst-34 fragments had the highest average Symbiodiniaceae cell densities (7.9e^5^ + 5.0e^5^ cells/cm^2^, Figure 2), inflating the average number of cell density for the worst performers (average for worst performers without Worst-34: 2.3e^5^ + 1.6e^5^ cells/cm^2^). Worst-2 and Worst-31 had significantly lower Symbiodiniaceae cell densities at the end of the experiment than other coral genets (Pairwise Wilcoxon rank sum test, Worst-2 lower than Worst-27 and Best-4, p=0.032; Worst-31 lower than Best-4, Best-12 and Worst-27, p=0.018, Supp. Mat. Table 1). Symbiont cell density was not significantly different among treatments (Kruskal-Wallis chi-squared=3.40, df=4, p=0.49), but based on fragment color change from the beginning to the end of the experiment, the heat treatment depressed Symbiodiniaceae traits significantly more than the combined stressor treatment (Pairwise Wilcoxon rank sum test, Heat lower than combined stressors, p=0.02). Buoyant growth rates were not significantly different among treatment conditions for genets that were dominated by similar symbiont communities (Best-4, Best-12, Best-20, Best-38, Worst-2 in Figure 1, Supp. Mat. Table 2).

**Figure 2.**
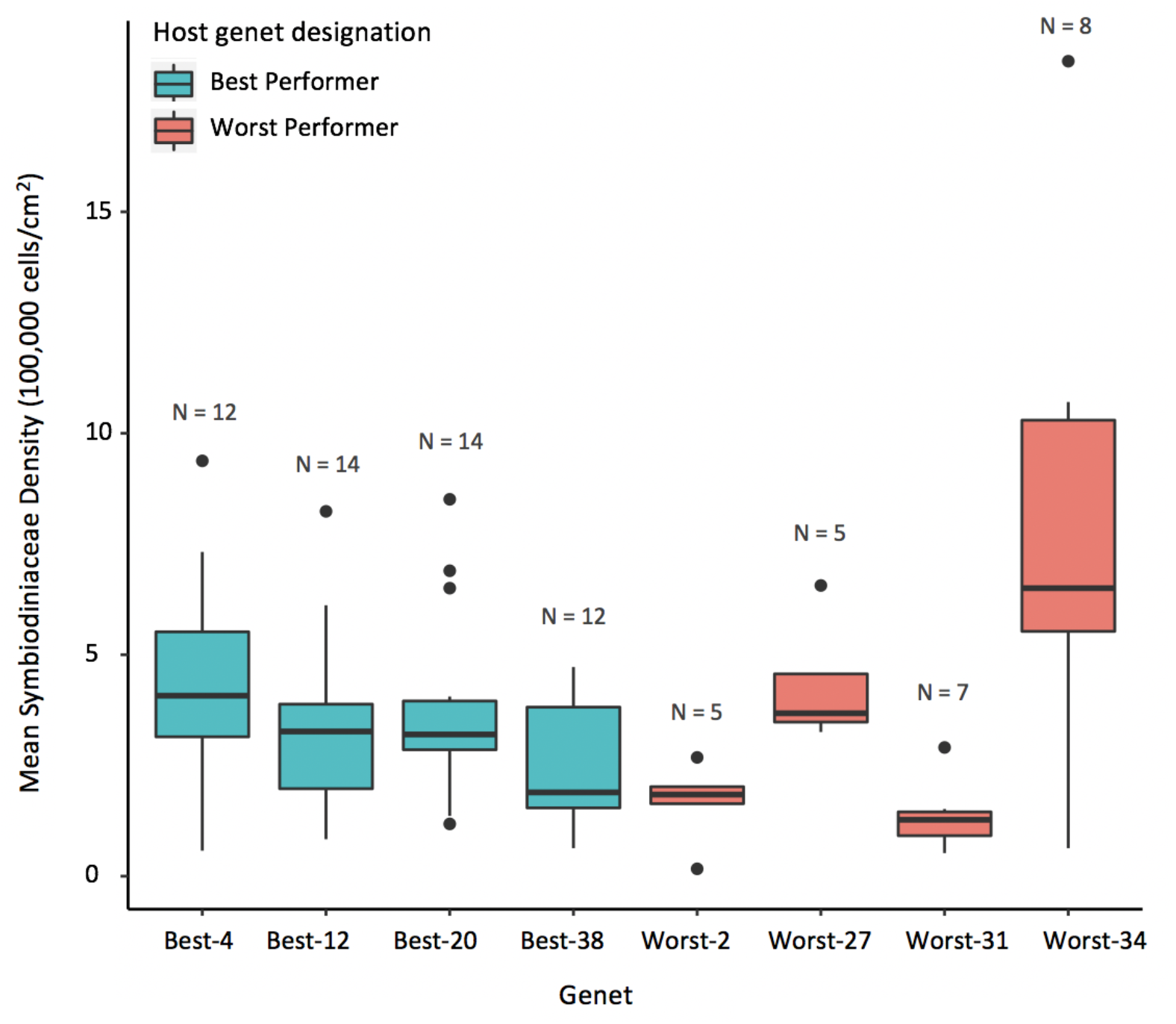
Average Symbiodiniaceae cell density counts from *Acropora millepora* genets sampled at the end of 10 days of experimental stress treatments. Sample sizes are provided above each genet. Worst-2 and Worst-31 had significantly lower Symbiodiniaceae cell density counts than other *A. millepora* genets (Worst-2 lower than Worst-27 and Best-4; Worst-31 lower than Best-4, Best-12, Best-20 and Worst-27; Supp. Mat. Table 1).

### Symbiodiniaceae community composition

Of the 102 fragments that survived the experimental conditions, 96 yielded sufficient DNA for amplicon sequencing (Figure 1). A total of 29,466,474 raw sequence reads were recovered from these samples. After processing through the DADA2 pipeline, paired reads per sample ranged from 589,944 to 996. Two samples that contained <10,000 reads (one Best-4 Control and one Best-4 Heat sample) were removed, leaving a total of 9,984,782 paired reads (Supp. Mat. Table 3) from which 232 ASVs were identified. The LULU pipeline resolved 12 ASVs from this dataset. One of these ASVs (represented by 12 reads total among two samples) was discarded because it did not produce any sequence similarities during the taxonomic assignment process. Some of the remaining 11 ASVs were subsequently assigned to the same Symbiodiniaceae species. Species assignment post-LULU ultimately identified a total of seven Symbiodiniaceae species, representing four genera, from the dataset: *Breviolum minutum, Cladocopium* 1232, *Cladocopium* 3k, *Durusdinium glynnii, Durusdinium trenchii, Durusdinium* 2, and *Gerakladium* 3.1 (Table 2). Fragments of Worst-31 collectively harbored the most Symbiodiniaceae diversity (all seven species), whereas fragments of Worst-2 collectively harbored the fewest (three) Symbiodiniaceae species (Table 2). Most individual coral fragments (94%, N = 88 of 94, Supp. Mat. Table 4) analyzed in this study harbored more than one Symbiodiniaceae species (see Table 2 for all detected species, Figure 1 for species comprising >0.1% of total Symbiodiniaceae community). Fragments in the *p*CO_2_ treatment collectively harbored the most (all seven) Symbiodiniaceae species, whereas fragments in the combined stressor treatment (bacterial addition, elevated temperature and *p*CO_2_) harbored the fewest (four) Symbiodiniaceae species (Table 2).

**Table 2.** Symbiodiniaceae amplicon sequence reads analyzed and Symbiodiniaceae species (or Internal Transcribed Spacer-2 region types) resolved from *Acropora millepora* fragments across several experimental stress treatments. The ‘Symbiodiniaceae species’ column lists the species or ITS-2 types cumulatively detected across all fragments of a host genet. C3k = *Cladocopium* 3k (NCBI Accession #: AY589737), C1232 = *Cladocopium* 1232 (Accession #: EU118163.1), D2 = *Durusdinium* 2 (Accession #: AY686649), and G3.1 = *Gerakladium* 3.1 (Accession #: KC597688).

| Host genet or treatment | # Reads | Symbiodiniaceae species |
| --- | --- | --- |
| **Best-4** | 1,586,758 | *Breviolum minutum*, C3k, C1232, *D. glynnii*, *Durisdinium trenchii* |
| **Best-12** | 1,198,509 | *B. minutum*, C3k, C1232, *D. glynnii* |
| **Best-20** | 2,293,707 | C3k, C1232, *D. glynnii*, *D. trenchii*, D2 |
| **Best-38** | 1,508,277 | C3k, C1232, *Durisdinium glynnii*, G3.1 |
| **Worst-2** | 639,548 | C3k, C1232, *D. glynnii* |
| **Worst-27** | 673,979 | C3k, C1232, *D. glynnii*, *D. trenchii* |
| **Worst-31** | 766,193 | *B. minutum*, C3k, C1232, *D. glynnii*, *D. trenchii*, D2, G3.1 |
| **Worst-34** | 1,317,811 | *B. minutum*, C3k, C1232, *D. glynnii* |
| **Control** | 2,288,947 | *B. minutum*, C3k, C1232, *D. glynnii*, *D. trenchii*, G3.1 |
| **Bacteria** | 1,764,362 | *B. minutum*, C3k, C1232, *D. glynnii*, *D. trenchii* |
| **Heat** | 1,900,830 | *B. minutum*, C3k, C1232, *D. glynnii*, *D. trenchii*, D2 |
| **pCO<sub>2</sub>** | 2,066,054 | *B. minutum*, C3k, C1232, *D. glynnii*, *D. trenchii*, D2, G3.1 |
| **Combined** | 1,964,589 | C3k, C1232, *D. glynnii*, *D. trenchii* |

### Coral genet, treatment, and Symbiodiniaceae community composition

Based on all ASVs identified by DADA2 (prior to processing the dataset through LULU), Symbiodiniaceae alpha diversity varied categorically by genet performance but did not differ based on treatment (Figure 3). Symbiont communities of worst performers were significantly more diverse by Shannon’s Diversity (H’) Index estimates (worst performer genets: 1.62, best performer genets: 1.20, Wilcoxon rank sum test: W = 2083, p < 0.001) and Simpson’s (1-D) Diversity estimates (worst performer genets: 0.68, best performer genets: 0.45, Wilcoxon rank sum test, W = 2118, p < 0.001). Shannon’s Diversity (H’) Index (Kruskal-Wallis chi-squared = 1.7423, df = 4, p = 0.78) and Simpson’s (1-D) Diversity results (Kruskal-Wallis chi-squared = 0.76442, df = 4, p = 0.94) for treatment were not significantly different. *A. millepora* Worst-27 was the most diverse genet (H’ = 1.84; 1-D = 0.76), whereas *A. millepora* Best-4 was the least diverse genet (H’ = 1.04; 1-D = 0.38; Figure 3).

**Figure 3.**
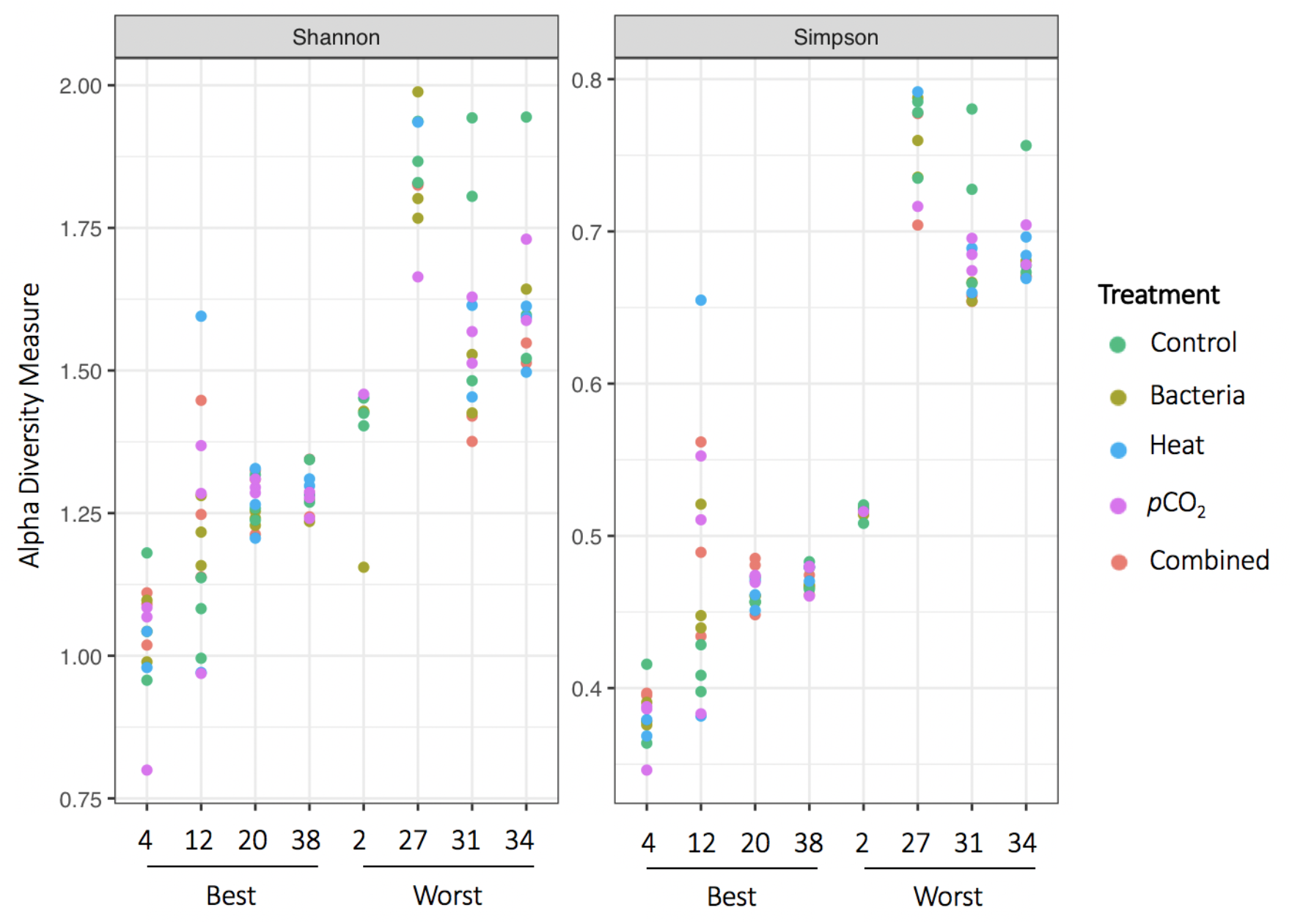
Alpha diversity of Symbiodiniaceae Internal Transcribed Spacer-2 (ITS-2) amplicon sequence variants by *Acropora millepora* genet and treatment. Shannon’s Diversity Index results are reported as H’, Simpson’s Diversity Index results are reported as 1-D. Alpha diversity indices were calculated from the full untrimmed dataset after processing by DADA2.

A permutation test for homogeneity of multivariate dispersions also indicated that variances, a proxy for beta diversity, were significantly lower among Symbiodininaceae communities associated with best performing *A. millepora* genets, than among communities associated with worst performing genets (df=1, p<0.001, Figure 4). In contrast, the same test showed no differences among the variances of Symbiodiniaceae communities exposed to stress versus control conditions (as a binary comparison, df=1, p=0.84), control conditions versus single stressors versus combined stressors (df=2, p=0.98), or among all of the different stress treatments (including combined stressors, df=4, p=0.96). Permutational multivariate analysis of variance supported that Symbiodiniaceae communities differed significantly between best versus worst performing *A. millepora* genets (Adonis with Bray-Curtis distance, p < 0.001, Figure 4). In contrast, Symbiodiniaceae communities among treatment conditions were not significantly different (Adonis with Bray-Curtis distance, p = 0.99). Within best or worst performing coral genets (as groups), stress treatment was also not associated with significantly different symbiont assemblages (Adonis with Bray-Curtis distance, Best performers, p = 0.75; Worst performers, p = 0.81).

**Figure 4.**
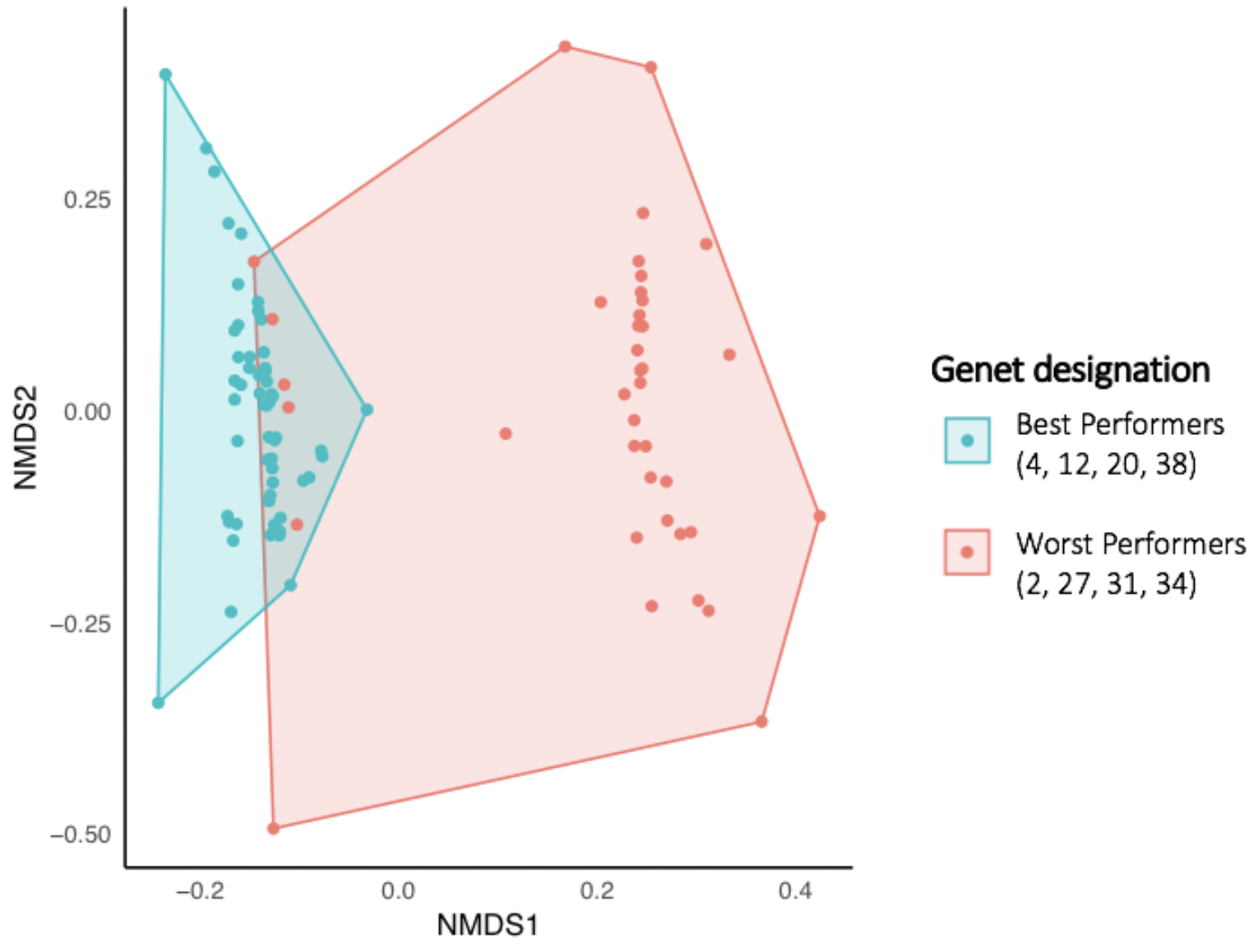
Symbiodiniaceae communities associated with *Acropora millepora* best performer genets differ from those associated with worst performer genets, based on non-metric multidimensional scaling (NMDS) using a Bray Curtis distance matrix. Variances, a proxy for beta diversity, are significantly higher among Symbiodininaceae communities associated with worst performer *A. millepora* genets, than among communities associated with best performer genets. The six Symbiodiniaceae communities with the lowest scores on NMDS1 represent the Worst-2 genet, collected from Davies Reef (along with Best-4).

When all three reef locations were incorporated into the regional GJAM model, it fitted the data well (Supp. Mat. Figure 2) and indicated that Symbiodiniaceae communities were sensitive to genet, but not to treatment (Figure 5 and sensitivity analysis shown in Supp. Mat. Figure 3). Using this regional model, we inversely predicted the *A. millepora* genet and experimental treatment given a particular Symbiodiniaceae community. Based on this process, some *A. millepora* genets were easily predicted; they had distinctive Symbiodiniaceae communities associated with them. This includes Best-4, Best-12, and Best-20, as well as Worst-2 (Figure 5). In contrast, prediction of experimental treatments from Symbiodiniaceae communities was not robust using the regional model (Figure 5). However, Symbiodiniaceae community was more sensitive to treatment in local GJAM models, which were run for Symbiodiniaceae communities at Pandora Reef and Rib Reef, respectively (Supp. Mat. Figure 3 compared to Supp. Mat. Table 5). In particular, treatment sensitivity values for Pandora Reef were higher than genet sensitivities for every stress type except bacterial addition and elevated temperature (Supp. Mat. Table 5).

**Figure 5.**
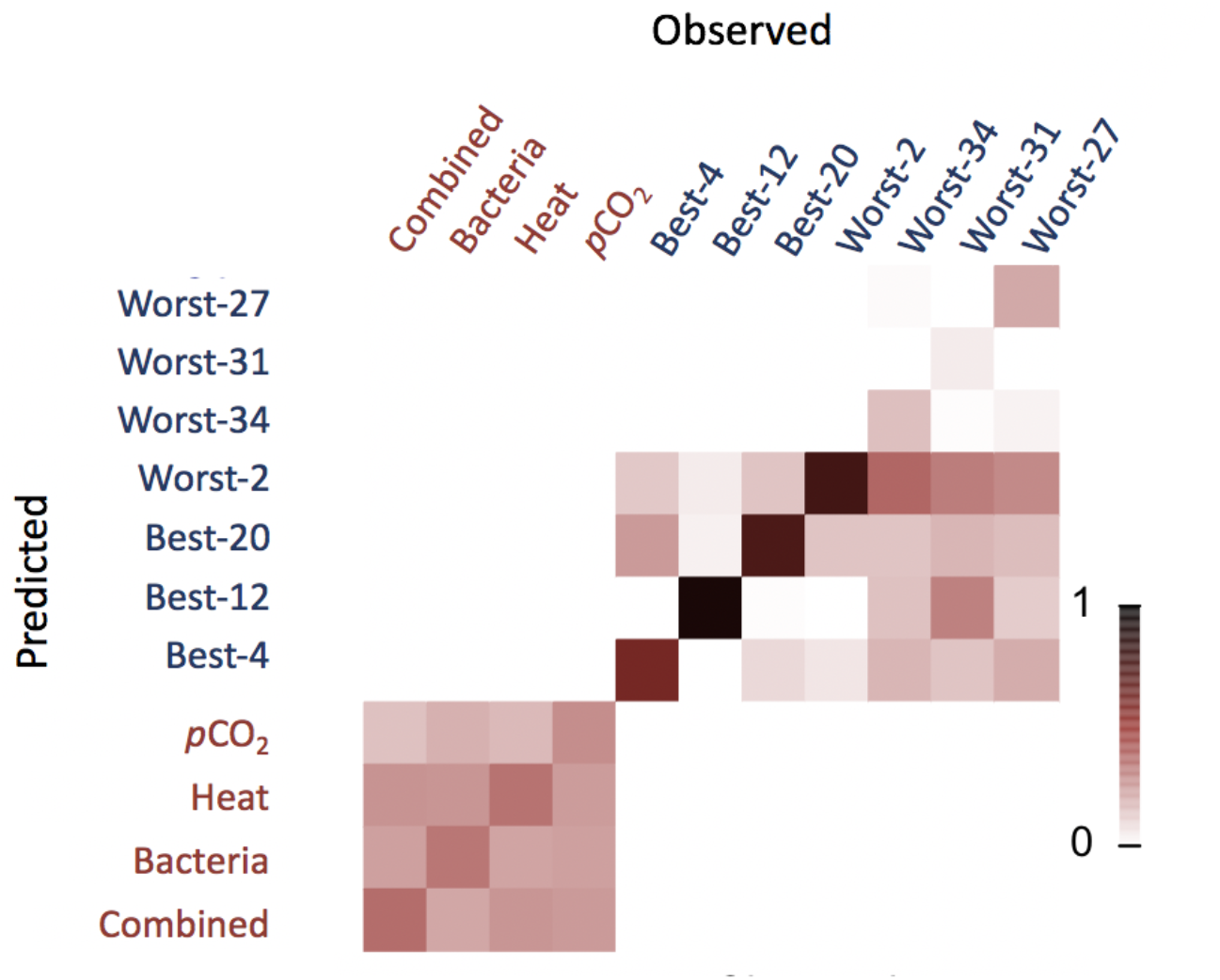
GJAM inverse predictions of *Acropora millepora* genet and experimental treatment based on Symbiodiniaceae community composition. Darker colors indicate that the *A. millepora* genet or treatment condition was more readily predicted from the Symbiodiniaceae community via the regional GJAM model.

### Symbiodiniaceae characteristics in best and worst performing coral holobionts

The dominant Symbiodiniaceae species in best performer *A. millepora* genets were more similar to each other than the dominant species associated with worst performer genet (Figure 1). Certain Symbiodiniaceae species were significantly associated with genet performance, or a stress treatment. Based on the indicspecies R package, *Cladocopium* 3k was a significant indicator species for best performer genets (p=0.005). *Cladocopium* 3k was also present at similar relative abundances within Worst-2 fragments, but this worst performer genet contained fewer background symbionts relative to best performer host genets (Table 2). *Cladocopium* 1232, *Durusdinium trenchii*, and *Durusdinium* 1 were indicator species for worst performer genets (p=0.005 for each). *Gerakladium* 3.1 was a significant indicator species for the*p*CO_2_ treatment (p=0.03) but only represented >1% of the total Symbiodiniaceae in one coral fragment in the study; there were no significant indicator species for the other treatments.

Results of the regional GJAM model indicated similar significant associations of Symbiodiniaceae species with certain coral genets (Figure 6). *Cladocopium* 3k_1 was strongly associated with Best-38 (labeled ‘intercept’ in Figure 6) and strongly disassociated with Worst-27, Worst-31, and Worst-34. *Cladocopium* 1232_1 was also strongly associated with Best-38, whereas *Cladocopium* 1232_2 and *Cladocopium* 3k_2 were strongly disassociated with that genet. *Cladocopium* 1232_3 was also strongly disassociated with Best-38 and strongly associated with Worst-27, Worst-31, and Worst-34 and Best-12. *Durusdinium trenchii* and *Durusdinium* 1 were strongly associated only with Worst-27 (Figure 6).

**Figure 6.**
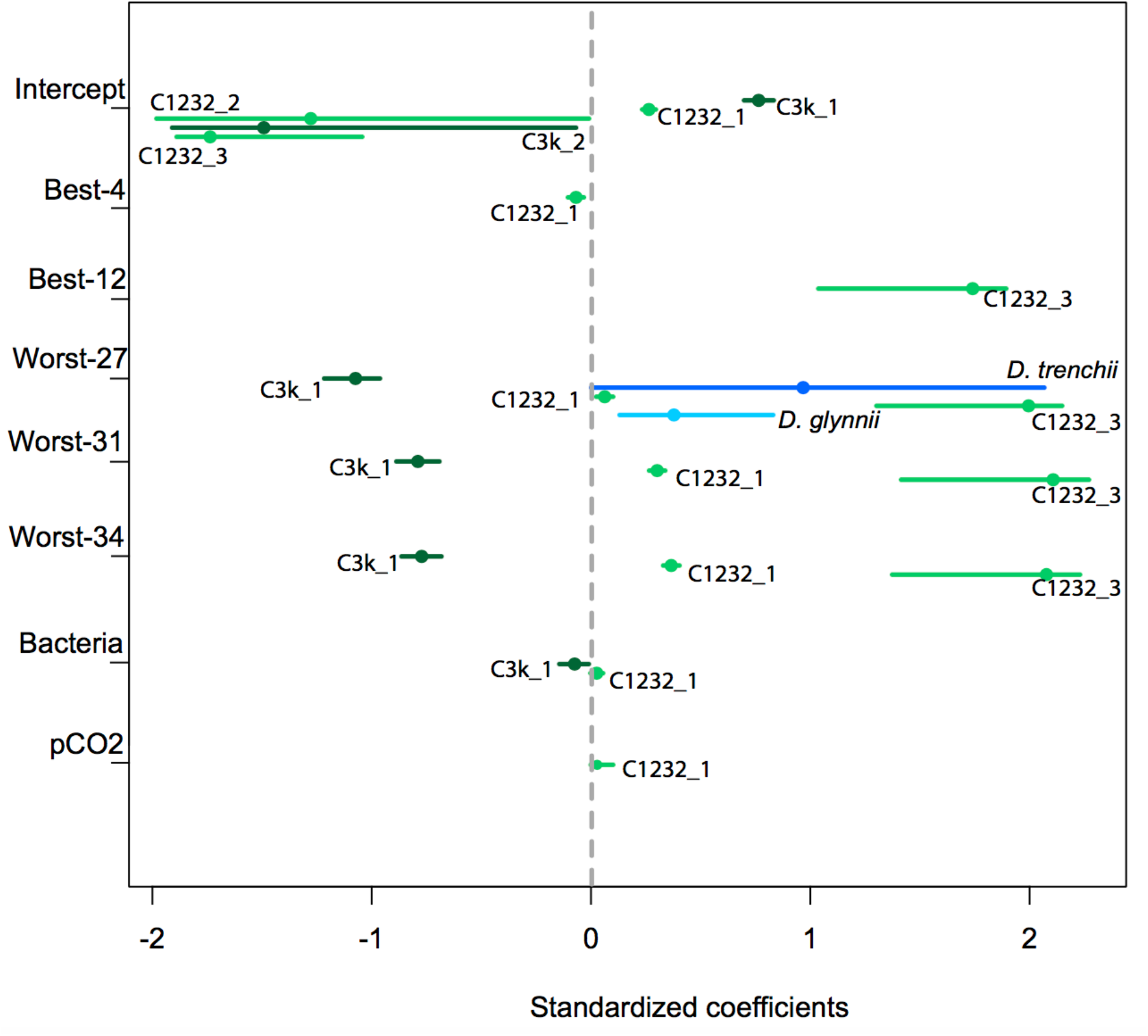
Posterior distributions of the significant effects of treatment and coral genet. Dots indicate the median and the segments expand to the 95% credible intervals. Light green = *Cladocopium* 3k ASVs; dark green = *Cladocopium* 1232 ASVs; light blue = *Durusdinium glynnii* ASVs; and dark blue = *Durusdinium trenchii* ASVs.

## Discussion

### Specific symbionts are associated with host survival under diverse stressors

This study demonstrates that different specific symbiont species are differentially associated with *A. millepora* host genets that exhibited high or low survivorship when challenged by various stressors. Both an indicator species analysis and a regional generalized joint attribute model (GJAM), identified a significant association between *Cladocopium* 3k and best performer genet(s) (Figure 6). Similar to this finding, Manzello et al., (2018) reported that 73% of the distribution of *Durusdinium trenchii* in a dominant reef-building Caribbean coral *(Orbicella faveolata)* was attributable to host identity. GJAM, which takes into account co-dependence among Symbiodininaceae species, also found a strong disassociation between *Cladocopium* 3k_1 and three of the four worst performer genets (Figure 6). *Cladocopium* 3k has previously been reported from a diversity of *Acropora* species (Barshis, Ladner, Oliver, & Palumbi, 2014; Smith, Pinzón, & LaJeunesse, 2009), including *A. millepora* (LaJeunesse et al., 2004). Most recently, *Cladocopium* 3k was reported from diverse host genera at mesophotic depths *(Echinophyllia, Fungia, Pachyseris*, and *Pavona* from 45-70m, Bongaerts et al., 2011) suggesting that this symbiont can cope with (or has locally adapted to) diverse light and temperature profiles. However, *Cladocopium 3k* has not previously been linked to stress tolerance; it is more often predicted to be thermally susceptible, for example, relative to *Durusdinium* 2 in American Samoan acroporids (e.g., Barshis et al., 2014). Based on recent coral transcriptomic studies, we infer that *Cladocopium* 3k may have contributed to the success of best performer host genets through differences in its baseline expression of various genes (relative to other Symbiodiniaceae species examined in this study), rather than transcriptional responses to stressors (Barshis et al., 2014; Leggat et al., 2011; H. M. Putnam, Mayfield, Fan, Chen, & Gates, 2013 but see Baumgarten et al., 2013). The physiology of *Cladocopium* 3k should be further characterized in monoculture and polyculture, as well as in the diverse Pacific *Acropora* coral species that host it. Beyond insights into symbiont physiology and function, such research efforts can yield insights into whether selection or complementarity effects are important in predicting and managing for holobiont stress tolerance.

We hypothesized that best performing host genets would contain higher relative abundances of stress-tolerant Symbiodiniaceae (e.g., *Durusinium trenchii)* than worst performing genets. Contrary to this, *Durusdinium glynnii* and *Durusdinium trenchii*, as well as *Cladocopium* 1232, were largely indicator species for worst performer genets based on indicator species analysis and GJAM. In Worst-27, *D. glynnii* and *D. trenchii* represented from 72 to 11% of the total Symbiodiniaceae community (Figure 1, Supp. Mat. Table 4). In all other coral fragments analyzed in this study, *Durusdinium* represented <1% of the total Symbiodiniaceae community (Supp. Mat. Table 4). Although some *Durusdinium* species have previously been associated with holobiont thermotolerance (e.g., the ITS-1 D in Berkelmans & van Oppen, 2006; Jones et al., 2008), *Durusdinium trenchii* is not necessarily tolerant of variable thermal conditions (Howells, Berkelmans, van Oppen, Willis, & Bay, 2013). Given the mortality suffered by Worst-27, there is no indication that *Durusdinium* species provided their host corals with tolerance to elevated temperature, *p*CO_2_, additions of *Vibrio owensii*, or combinations thereof in this experiment (see mortality and relative abundance of *Durusdinium* species in Figure 1). The significant associations among symbiont species and host genets, but not treatments, at the regional scale (Figure 5) suggests that Symbiodininaceae communities might be more strongly structured by the host (or biotic interactions) in *A. millepora*, rather than imposed by the environment. Yet, within a reef, where there is less variation in host genet, some influence of environment (i.e., treatment) on symbiont community composition becomes apparent (Supp. Mat. Table 5). This fits the general expectation that history and evolutionary processes, in this case contributing to a heterogeneous distribution of host genets, might have a greater impact at regional (among reefs) as opposed to local (within reef) scales (e.g., Witman, Etter, & Smith, 2004).

### Host genet, but not stress type, is predictive of symbiont community attributes among reefs

Beyond individual Symbiodiniaceae species, symbiont community attributes (alpha and beta diversity) also differed significantly between best versus worst performer coral genets, but not stress type. Best performer host genets had fewer Symbiodiniaceae ASVs and higher community evenness than worst performer host genets (Figure 3). Symbiont communities among best performer coral fragments were similar to each other, whereas symbiont communities among fragments of worst performer coral genets exhibited high variance or beta diversity (Figure 4). These findings agree with a recent general observation that dysbiotic host individuals vary more in microbial community composition than healthy host individuals – the ‘Anna Karenina Principle’ or AKP (Zaneveld et al., 2017). This has previously been documented in experimentally stressed coral-associated bacterial communities (Zaneveld et al., 2016) and in the survival of coral juveniles from distinct parental genets (Quigley et al., 2016). Inverse predictions of host genet from Symbiodiniaceae community composition by the regional GJAM model support this: the model easily predicted most of the best performer coral genets (Best-4, Best-12, Best-20 in Figure 5), but could only reliably predict one worst performer genet (Worst-2). It should be noted, however, that coral fragments exposed to stress treatments did not exhibit higher variance in symbiont communities compared to control fragments at the end of the experiment. Thus, our data fit with the AKP in that healthy (best performer) coral genets host more similar Symbiodiniaceae communities, but our data do not represent the AKP in terms of exhibiting a “constrained ‘core’ of control microbiomes surrounded by a large ‘halo’ or ‘smear’ of stressed or diseased microbiomes”, as described by Zaneveld et al., (2017). This is the first study to directly assess whether the AKP is applicable to Symbiodiniaceae communities. These symbionts join a growing list of microbial communities, particularly those associated with various organ systems in primates (Chen et al., 2015; Halfvarson et al., 2017; Moeller et al., 2013; Wu et al., 2016) that exhibit stochastic responses to environmental stressors. Future HTS studies examining Symbiodiniaceae diversity from a whole community perspective can further clarify the relationship between microbiome stability and host performance as reefs and other systems continue to be challenged with diverse stressors.

### Absence of tradeoffs in symbiont community under challenge by diverse stressors

Fragments of best performer coral genets were dominated by *Cladocopium* 3k (with a lower relative abundance of *Cladocopium* 1232) regardless of the stress treatment that fragments experienced in this study (Figure 1). Symbiodiniaceae community composition also did not change significantly among treatments within a given worst performer genet (Figure 1). Thus, we reject our hypothesis that fragments of a given host genet exposed to different stress treatments would differ significantly in their symbiont community composition and diversity, as well as from that of control fragments. This is consistent with recent findings that bacterial communities of *Acropora tenuis* are highly specific to host genet regardless of challenge with various environmental stressors (Glasl, Smith, Bourne, & Webster, 2019). Furthermore, Symbiodiniaceae communities were minimally different in fragments experiencing multiple stressors, as compared to one stressor (Figure 1), although the total diversity of Symbiodiniaceae species (including those present at <1% of the total community) was reduced in fragments experiencing combined stressors (Table 2).

Tradeoffs might have been expected in the growth rates of hosts harboring different Symbiodiniaceae communities under control versus stress treatments. For example, it has previously been documented that corals dominated by some symbionts in *Durusdinium* exhibit higher thermotolerance, but slower growth rates, than the same host species dominated by some symbionts in *Cladocopium* (Jones & Berkelmans, 2010; Little et al., 2004). In this study, symbiont community compositions were determined post-hoc and unfortunately, sample sizes proved insufficient to test for tradeoffs among distinct symbiont communities. However, five host genets (Best-4, Best-12, Best-20, Best-38 and Worst-2) contained a particular symbiont community (~70% *Cladocopium* 3k and nearly 30% *Cladocopium* 1232 + background symbionts, Figure 1, Table 2). Across these fragments dominated by *Cladocopium* 3k (and to a lesser extent, 1232), there was no apparent change in host growth rate under control versus stress conditions, or among different stressors, based on log growth of fragments over the 10-day duration of this study (Supp. Mat. Table 2). The lack of detectable tradeoffs in hosting the same Symbiodiniaceae community under different stressors found in this study agrees with findings by Wright et al. *(submitted)* for host genets. Based on measurements of seven host-specific traits (e.g., host chromoprotein content, instant calcification rate, buoyant weight increase) from the same coral fragments (as well as 30 additional host genets), host genets that performed well under one stress tended to survive other stressors as well (Wright et al. *submitted).* Thus, our findings related to symbiont communities are not likely to be mere artifacts of the eight host genets examined here.

### Methodological considerations

This study compares Symbiodiniaceae communities in fragments of individual coral genets that experienced stress treatments for 10 days to fragments that experienced ambient conditions during the experimental period. Data on symbiont community composition at T0 are not available and therefore we cannot directly assess the extent to which symbiont communities were heterogeneously distributed in colonies prior to fragmenting for the experiment, or whether stress treatments caused symbiont communities to shift within individual coral fragments. Specifically, *Cladocopium 3k* could have been lost from stress-treated fragments of Worst-31 over the course of the experiment, given the symbiont community composition in control Worst-31 fragments at the final time point (Figure 1). However, the dominant symbiont species present in the three control fragments of each coral genet at the final time point were remarkably similar (Figure 1) suggesting that intracolony spatial heterogeneity in symbiont distributions and symbiont shuffling have minimal explanatory power (if any) in this dataset.

In this study, genets were generally assessed in terms of their performance, yet our characterization of the symbiont community compositions and cell densities of worst performer genets was based on surviving fragments only. We cannot examine post-hoc whether individual worst performer fragments that experienced mortality had unique symbiont community characteristics (relative to surviving worst performer fragments of the same genet) that ultimately contributed to fragment death. Quantification of symbiont density per host genet in this study is also based on surviving fragments only. Many worst performer coral fragments bleached as they died, but these fragments were not represented in average symbiont density calculations for worst performer genets. Thus, the extent to which stressors negatively impacted worst performer genet health, and the difference in bleaching that occurred between worst versus best performer host genets, is likely underestimated in this study.

This study is unique in that symbiont community composition was examined from host genets that were known best or worst performers, based on their survival under various stressful conditions. Importantly, we found that symbiont community composition and diversity metrics were highly similar among best performer host genets, whereas symbiont compositions were more ‘dysbiotic’ or variable in worst performer genets. Indicator species analysis and the regional GJAM model also highlighted several Symbiodiniaceae species that were strongly associated with best or worst performer genets. The presence of a similar symbiont community across diverse stressors in best performer host genets provides (limited) hope for coral reefs: up to a point, best performer host-symbiont combinations can potentially resist diverse or multiple stressors. The extent to which we can identify and promote specific symbionts (e.g., *Cladocopium* 3k), communities or their characteristics (e.g., target levels of alpha, beta diversity) across diverse host genets and species is a logical next goal in managing for coral reef resistance and resilience.

## Acknowledgments

We thank Noah Workman at Georgia Genomics Facility for consultation regarding Illumina sequencing and Avery Zaleski for assistance processing sequence data. This study was made possible through support from the U.S. National Science Foundation (OCE #1635798 to AMSC), a Sigma Xi Grant-in-Aid of Research to LHK, and an AIMS funding to LKB. Samples were collected under general AIMS permit G11/34671.1 and G14/37318.1. The authors would like to acknowledge experimental assistance by Maria Nayfa and Ari Muskat and have no conflicts of interest to report.

## References

Anderson, M. J., & Walsh, D. C. I. (2013). PERMANOVA, ANOSIM, and the Mantel test in the face of heterogeneous dispersions: What null hypothesis are you testing? Ecological Monographs, 83(4), 557–574. https://doi.org/10.1890/12-2010.1

Baker, A. C. (2001). Reef corals bleach to survive change. Nature, 411, 765–766. https://doi.org/10.1038/35081151

Baker, A. C., Starger, C. J., McClanahan, T. R., & Glynn, P. W. (2004). Corals’ adaptive response to climate change. Nature, 430, 741. https://doi.org/10.1038/430741a

Barshis, D. J., Ladner, J. T., Oliver, T. A., & Palumbi, S. R. (2014). Lineage-specific transcriptional profiles of *Symbiodinium* spp. unaltered by heat stress in a coral host. Molecular Biology and Evolution, 31(6), 1343–1352. https://doi.org/10.1093/molbev/msu107

Baumgarten, S., Bayer, T., Aranda, M., Liew, Y. J., Carr, A., Micklem, G., & Voolstra, C. R. (2013). Integrating microRNA and mRNA expression profiling in Symbiodinium microadriaticum, a dinoflagellate symbiont of reef-building corals. BMC Genomics, 14, 704. https://doi.org/10.1186/1471-2164-14-704

Bay, L. K., Doyle, J., Logan, M., & Berkelmans, R. (2016). Recovery from bleaching is mediated by threshold densities of background thermo-tolerant symbiont types in a reef-building coral. Royal Society Open Science, 3, 160322. https://doi.org/10.1098/rsos.160322

Berkelmans, R., & van Oppen, M. J. H. (2006). The role of zooxanthellae in the thermal tolerance of corals: A “nugget of hope” for coral reefs in an era of climate change. Proceedings of the Royal Society B: Biological Sciences, 273, 2305–2312. https://doi.org/10.1098/rspb.2006.3567

Bongaerts, P., Sampayo, E. M., Bridge, T. C. L., Ridgway, T., Vermeulen, F., Englebert, N., … Hoegh-Guldberg, O. (2011). *Symbiodinium* diversity in mesophotic coral communities on the Great Barrier Reef: a first assessment. Marine Ecology Progress Series, 439, 117–126. https://doi.org/10.3354/meps09315

Brading, P., Warner, M. E., Davey, P., Smith, D. J., Achterberg, E. P., & Suggett, D. J. (2011). Differential effects of ocean acidification on growth and photosynthesis among phylotypes of Symbiodinium (Dinophyceae). Limnology and Oceanography, 56(3), 927–938. https://doi.org/10.4319/lo.2011.56.3.0927

Brener-Raffalli, K., Clerissi, C., Vidal-Dupiol, J., Adjeroud, M., Bonhomme, F., Pratlong, M., … Toulza, E. (2018). Thermal regime and host clade, rather than geography, drive Symbiodinium and bacterial assemblages in the scleractinian coral Pocillopora damicornis sensu lato. Microbiome, 6, 39. https://doi.org/10.1186/s40168-018-0423-6

Callahan, B. J., McMurdie, P. J., Rosen, M. J., Han, A. W., Johnson, A. J. A., & Holmes, S. P. (2016). DADA2: High-resolution sample inference from Illumina amplicon data. Nature Methods, 13, 581–583. https://doi.org/10.1038/nmeth.3869

Cantin, N. E., van Oppen, M. J. H., Willis, B. L., Mieog, J. C., & Negri, A. P. (2009). Juvenile corals can acquire more carbon from high-performance algal symbionts. Coral Reefs, 28, 405–414. https://doi.org/10.1007/s00338-009-0478-8

Chen, Y., Guo, J., Qian, G., Fang, D., Shi, D., Guo, L., & Li, L. (2015). Gut dysbiosis in acute-on-chronic liver failure and its predictive value for mortality. Journal of Gastroenterology and Hepatology, 30(9), 1429–1437. https://doi.org/10.1111/jgh.12932

Clark, J. S., Nemergut, D., Seyednasrollah, B., Turner, P. J., & Zhang, S. (2017). Generalized joint attribute modeling for biodiversity analysis: Median-zero, multivariate, multifarious data. Ecological Monographs, 87(1), 34–56. https://doi.org/10.1002/ecm.1241

Cunning, R., Gates, R. D., & Edmunds, P. J. (2017). Using high-throughput sequencing of ITS2 to describe *Symbiodinium* metacommunities in St. John, US Virgin Islands. PeerJ, 5, e3472. https://doi.org/10.7717/peerj.3472

De Cáceres, M., & Legendre, P. (2009). Associations between species and groups of sites: indices and statistical inference. Ecology, 90(12), 3566–3574. Retrieved from http://adn.biol.umontreal.ca/~numericalecology/Reprints/De_Caceres_&_Legendre_Ecology_2009.pdf

Fox, J. W. (2005). Interpreting the “selection effect” of biodiversity on ecosystem function. Ecology Letters, 8, 846–856. https://doi.org/10.1111/j.1461-0248.2005.00795.x

Frøslev, T. G., Kjøller, R., Bruun, H. H., Ejrnæs, R., Brunbjerg, A. K., Pietroni, C., & Hansen, A. J. (2017). Algorithm for post-clustering curation of DNA amplicon data yields reliable biodiversity estimates. Nature Communications, 8(1). https://doi.org/10.1038/s41467-017-01312-x

Glasl, B., Smith, C. E., Bourne, D. G., & Webster, N. S. (2019). Disentangling the effect of host-genotype and environment on the microbiome of the coral Acropora tenuis. PeerJ, 7, e6377. https://doi.org/10.7717/peerj.6377

Green, E. A., Davies, S. W., Matz, M. V., & Medina, M. (2014). Quantifying cryptic Symbiodinium diversity within Orbicella faveolata and Orbicella franksi at the Flower Garden Banks, Gulf of Mexico. PeerJ, 2, e386. https://doi.org/10.7717/peerj.386

Halfvarson, J., Brislawn, C. J., Lamendella, R., Vázquez-Baeza, Y., Walters, W. A., Bramer, L. M., … Jansson, J. K. (2017). Dynamics of the human gut microbiome in inflammatory bowel disease. Nature Microbiology, 2, 17004. https://doi.org/10.1038/nmicrobiol.2017.4

Hauff, B., Cervino, J. M., Haslun, J. A., Krucher, N., Wier, A. M., Mannix, A. L., … Strychar, K. B. (2014). Genetically divergent *Symbiodinium* sp. display distinct molecular responses to pathogenic *Vibrio* and thermal stress. Diseases of Aquatic Organisms, 112, 149–159. https://doi.org/10.3354/dao02802

Howells, E. J., Berkelmans, R., van Oppen, M. J. H., Willis, B. L., & Bay, L. K. (2013). Historical thermal regimes define limits to coral acclimatization. Ecology, 94(5), 1078–1088.

Hughes, T. P., Kerry, J. T., Álvarez-Noriega, M., Álvarez-Romero, J. G., Anderson, K. D., Baird, A. H., … Wilson, S. K. (2017). Global warming and recurrent mass bleaching of corals. Nature, 543, 373–377. https://doi.org/10.1038/nature21707

Hughes, T. P., Kerry, J. T., Baird, A. H., Connolly, S. R., Dietzel, A., Eakin, C. M., … Torda, G. (2018). Global warming transforms coral reef assemblages. Nature, 556, 492–496. https://doi.org/10.1038/s41586-018-0041-2

Hume, B. C. C., Ziegler, M., Poulain, J., Pochon, X., Romac, S., Boissin, E., … Voolstra, C. R. (2018). An improved primer set and amplification protocol with increased specificity and sensitivity targeting the *Symbiodinium* ITS2 region. PeerJ, 6, e4816. https://doi.org/10.7717/peerj.4816

Jones, A., & Berkelmans, R. (2010). Potential costs of acclimatization to a warmer climate: growth of a reef coral with heat tolerant vs. sensitive symbiont types. PLoS ONE, 5(5), e10437. https://doi.org/10.1371/journal.pone.0010437

Jones, A. M., Berkelmans, R., van Oppen, M. J. H., Mieog, J. C., & Sinclair, W. (2008). A community change in the algal endosymbionts of a scleractinian coral following a natural bleaching event: Field evidence of acclimatization. Proceedings of the Royal Society B: Biological Sciences, 275, 1359–1365. https://doi.org/10.1098/rspb.2008.0069

Kaniewska, P., Campbell, P. R., Kline, D. I., Rodriguez-Lanetty, M., Miller, D. J., Dove, S., & Hoegh-Guldberg, O. (2012). Major cellular and physiological impacts of ocean acidification on a reef building coral. PLoS ONE, 7(4), e34659. https://doi.org/10.1371/journal.pone.0034659

Kenkel, C. D., & Bay, L. K. (2018). Exploring mechanisms that affect coral cooperation: symbiont transmission mode, cell density and community composition. PeerJ, 6, e6047. https://doi.org/10.7717/peerj.6047

LaJeunesse, T. C., Bhagooli, R., Hidaka, M., DeVantier, L., Done, T., Schmidt, G. W., … Hoegh-Guldberg, O. (2004). Closely related *Symbiodinium* spp. differ in relative dominance in coral reef host communities across environmental, latitudinal and biogeographic gradients. Marine Ecology Progress Series, 284, 147–161. https://doi.org/10.3354/meps284147

LaJeunesse, T. C., Pettay, D. T., Sampayo, E. M., Phongsuwan, N., Brown, B., Obura, D. O., … Fitt, W. K. (2010). Long-standing environmental conditions, geographic isolation and host—symbiont specificity influence the relative ecological dominance and genetic diversification of coral endosymbionts in the genus *Symbiodinium*. Journal of Biogeography, 37, 785–800. https://doi.org/10.1111/j.1365-2699.2010.02273.x

LaJeunesse, T. C., Smith, R. T., Finney, J., & Oxenford, H. (2009). Outbreak and persistence of opportunistic symbiotic dinoflagellates during the 2005 Caribbean mass coral “bleaching” event. Proceedings of the Royal Society B: Biological Sciences, 276(1676), 4139–4148. https://doi.org/10.1098/rspb.2009.1405

Lee, M. J., Jeong, H. J., Jang, S. H., Lee, S. Y., Kang, N. S., Lee, K. H., … LaJeunesse, T. C. (2016). Most Low-Abundance “Background” *Symbiodinium* spp. Are Transitory and Have Minimal Functional Significance for Symbiotic Corals. Microbial Ecology, 71(3), 771–783. https://doi.org/10.1007/s00248-015-0724-2

Leggat, W., Seneca, F., Wasmund, K., Ukani, L., Yellowlees, D., & Ainsworth, T. D. (2011). Differential Responses of the Coral Host and Their Algal Symbiont to Thermal Stress. PLoS ONE, 6(10), e26687. https://doi.org/10.1371/journal.pone.0026687

Little, A. F., van Oppen, M. J. H., & Willis, B. L. (2004). Flexibility in Algal Endosymbioses Shapes Growth in Reef Corals. Science, 304(5676), 1492–1495. https://doi.org/10.1126/science.1095733

Loreau, M., Naeem, S., Inchausti, P., Bengtsson, J., Grime, J. P., Hector, A., … Wardle, D. A. (2001). Biodiversity and Ecosystem Functioning : Current Knowledge and Future Challenges. Science, 294(5543), 804–809. https://doi.org/10.1126/science.1064088

Lundgren, P., Vera, J. C., Peplow, L., Manel, S., & van Oppen, M. J. H. (2013). Genotype – environment correlations in corals from the Great Barrier Reef. BMC Genetics, 14, 9. https://doi.org/10.1186/1471-2156-14-9

Manzello, D. P., Matz, M. V., Enochs, I. C., Valentino, L., Carlton, R. D., Kolodziej, G., … Jankulak, M. (2018). Role of host genetics and heat tolerant algal symbionts in sustaining populations of the endangered coral Orbicella faveolata in the Florida Keys with ocean warming. Global Change Biology, 25, 1016–1031. https://doi.org/10.1111/gcb.14545

Mieog, J. C., Olsen, J. L., Berkelmans, R., Bleuler-Martinez, S. A., Willis, B. L., & van Oppen, M. J. H. (2009). The roles and interactions of symbiont, host and environment in defining coral fitness. PLoS ONE, 4(7), e6364. https://doi.org/10.1371/journal.pone.0006364

Moeller, A. H., Shilts, M., Li, Y., Rudicell, R. S., Lonsdorf, E. V., Pusey, A. E., … Ochman, H. (2013). Siv-induced instability of the chimpanzee gut microbiome. Cell Host and Microbe, 14, 340–345. https://doi.org/10.1016/j.chom.2013.08.005

Putnam, H. M., Mayfield, A. B., Fan, T. Y., Chen, C. S., & Gates, R. D. (2013). The physiological and molecular responses of larvae from the reef-building coral *Pocillopora damicornis* exposed to near-future increases in temperature and pCO2. Marine Biology, 160, 2157–2173. https://doi.org/10.1007/s00227-012-2129-9

Putnam, H. M., Stat, M., Pochon, X., & Gates, R. D. (2012). Endosymbiotic flexibility associates with environmental sensitivity in scleractinian corals. Proceedings of the Royal Society B: Biological Sciences, 279, 4352–4361. https://doi.org/10.1098/rspb.2012.1454

Quigley, K. M., Bay, L. K., & Willis, B. L. (2017). Temperature and Water Quality-Related Patterns in Sediment-Associated *Symbiodinium* Communities Impact Symbiont Uptake and Fitness of Juveniles in the Genus Acropora. Frontiers in Marine Science, 4, 401. https://doi.org/10.3389/fmars.2017.00401

Quigley, K. M., Davies, S. W., Kenkel, C. D., Willis, B. L., Matz, M. V., & Bay, L. K. (2014). Deep-sequencing method for quantifying background abundances of Symbiodinium types: Exploring the rare Symbiodinium biosphere in reef-building corals. PLoS ONE, 9(4), e94297. https://doi.org/10.1371/journal.pone.0094297

Quigley, K. M., Warner, P. A., Bay, L. K., & Willis, B. L. (2018). Unexpected mixed-mode transmission and moderate genetic regulation of *Symbiodinium* communities in a brooding coral. Heredity, 121, 524–536. https://doi.org/10.1038/s41437-018-0059-0

Quigley, K. M., Willis, B. L., & Bay, L. K. (2016). Maternal effects and Symbiodinium community composition drive differential patterns in juvenile survival in the coral Acropora tenuis. Royal Society Open Science, 3(10), 160471. https://doi.org/10.1098/rsos.160471

Quigley, K. M., Willis, B. L., & Bay, L. K. (2017). Heritability of the Symbiodinium community in vertically- and horizontally-transmitting broadcast spawning corals. Scientific Reports, 7, 8219. https://doi.org/10.1038/s41598-017-08179-4

Rouzé, H., Lecellier, G., Saulnier, D., & Berteaux-Lecellier, V. (2016). Symbiodinium clades A and D differentially predispose Acropora cytherea to disease and Vibrio spp. colonization. Ecology and Evolution, 6(2), 560–572. https://doi.org/10.1002/ece3.1895

Silverstein, R. N., Cunning, R., & Baker, A. C. (2015). Change in algal symbiont communities after bleaching, not prior heat exposure, increases heat tolerance of reef corals. Global Change Biology, 21, 236–249. https://doi.org/10.1111/gcb.12706

Silverstein, R. N., Cunning, R., & Baker, A. C. (2017). Tenacious D: Symbiodinium in clade D remain in reef corals at both high and low temperature extremes despite impairment. The Journal of Experimental Biology, 220, 1192–1196. https://doi.org/10.1242/jeb.148239

Smith, R. T., Pinzón, J. H., & Lajeunesse, T. C. (2009). Symbiodinium (dinophyta) diversity and stability in aquarium corals. Journal of Phycology, 45, 1030–1036. https://doi.org/10.1111/j.1529-8817.2009.00730.x

Stimson, J., & Kinzie, R. A. (1991). The temporal pattern and rate of release of zooxanthellae from the reef coral *Pocillopora damicornis* (Linnaeus) under nitrogen-enrichment and control conditions. Journal of Experimental Marine Biology and Ecology, 153, 63–74. https://doi.org/10.1016/S0022-0981(05)80006-1

van Oppen, M. J. H., Bongaerts, P., Frade, P., Peplow, L. M., Boyd, S. E., Nim, H. T., & Bay, L. K. (2018). Adaptation to reef habitats through selection on the coral animal and its associated microbiome. Molecular Ecology, 27, 2956–2971. https://doi.org/10.1111/mec.14763

Witman, J. D., Etter, R. J., & Smith, F. (2004). The relationship between regional and local species diversity in marine benthic communities: A global perspective. Proceedings of the National Academy of Sciences, 101(44), 15664–15669. https://doi.org/10.1073/pnas.0404300101

Wu, J., Peters, B. A., Dominianni, C., Zhang, Y., Pei, Z., Yang, L., … Ahn, J. (2016). Cigarette smoking and the oral microbiome in a large study of American adults. ISME Journal, 10, 2435–2446. https://doi.org/10.1038/ismej.2016.37

Zaneveld, J. R., Burkepile, D. E., Shantz, A. A., Pritchard, C. E., McMinds, R., Payet, J. P., … Thurber, R. V. (2016). Overfishing and nutrient pollution interact with temperature to disrupt coral reefs down to microbial scales. Nature Communications, 7, 11833. https://doi.org/10.1038/ncomms11833

Zaneveld, J. R., McMinds, R., & Thurber, R. V. (2017). Stress and stability: Applying the Anna Karenina principle to animal microbiomes. Nature Microbiology, 2, 17121. https://doi.org/10.1038/nmicrobiol.2017.121

Ziegler, M., Arif, C., Burt, J. A., Dobretsov, S., Roder, C., LaJeunesse, T. C., & Voolstra, C. R. (2017). Biogeography and molecular diversity of coral symbionts in the genus *Symbiodinium* around the Arabian Peninsula. Journal of Biogeography, 44(3), 674–686. https://doi.org/10.1111/jbi.12913

Ziegler, M., Eguíluz, V. M., Duarte, C. M., & Voolstra, C. R. (2018). Rare symbionts may contribute to the resilience of coral-algal assemblages. ISME Journal, 12, 161–172. https://doi.org/10.1038/ismej.2017.151

Ziegler, M., Stone, E., Colman, D., Takacs-Vesbach, C., & Shepherd, U. (2018). Patterns of Symbiodinium (Dinophyceae) diversity and assemblages among diverse hosts and the coral reef environment of Lizard Island, Australia. Journal of Phycology, 54(4), 447–460. https://doi.org/10.1111/jpy.12749

