## Supplementary Materials for "Symbiont community diversity is more constrained in holobionts that tolerate diverse stressors"

**Supplementary Material**

**Methods**

*Study species and experimental design*

On October 20th, 2014 each *Acropora millepora* coral genet was fragmented into replicate branches, which were mounted on aragonite plugs using super glue and placed in sequential order on trays. Trays were maintained in six indoor holding tanks which were supplied with 0.2uM filtered seawater at 27°C. Three (AI Aqua Illumination) lights were suspended above each tank, providing an average underwater light intensity of 80 umol photons m^-2^ s^-1^ on a 10:14 hour light-dark cycle. Corals were fed twice daily with freshly hatched Artemia nauplii and cleaned three times per week to prevent algal overgrowth. Coral fragments were acclimatized to these common garden conditions for approximately five months (128 days for genets #2-34 and 82 days for genet #38).

On March 2nd, 2015, coral fragments were placed into 50-L treatment tanks fitted with 3.5-watt Turbelle nanostream 6015 pumps (Tunze, Germany) with flow-through seawater (~25 liters/hour) at the same temperature and light conditions as in the large holding tanks. From March 6 - March 13th, temperature and pCO_2_ were incrementally ramped in their respective tanks (N=3 per treatment) to achieve the experimental treatment levels. Beginning on March 12th, coral fragments were photographed daily to quantify bleaching and lesion progression. Any fragments exhibiting significant tissue sloughing were removed from aquaria, buoyant weighed, snap frozen in liquid nitrogen and time of death was recorded. Treatment continued until ~25% mortality was observed in any treatment. Surviving coral fragments were photographed, buoyant weighed, and snap frozen in liquid nitrogen on March 22nd.

Coral fragments were stored at -80°C until processing. Tissue was removed from coral skeletons using an airbrush and homogenized for 60 seconds using a Pro250 homogenizer (Perth Scientific Equipment, AUS). A 250μl aliquot of the tissue homogenate was fixed with 5% formalin in FSW and used to quantify Symbiodiniaceae cell density. The remaining homogenate was centrifuged for 2 min at 3500 rcf to separate host and symbiont fractions for molecular work. The Symbiodiniaceae cell pellet was washed with FSW, resuspended in 2ml FSW and both fractions were frozen in 96-well tissue culture plates at -80°C. Coral skeletons were rinsed with 5% bleach then dried at room temperature. Skeletal surface area was quantified using the single wax dipping method and skeletal volume was determined by calculating water displacement in a graduated cylinder.

**Figure 1. Map indicating the source reefs for the eight *Acropora millepora* genets analyzed in this study.** Distinct coral genets were collected from Pandora Island (Worst-27, Worst-31, Worst-34), Rib Reef (Best-12, Best-20, Best-38), and Davies Reef lagoon (Best-4, Worst-2).

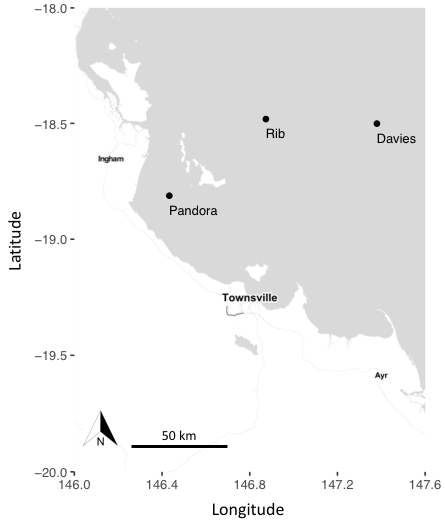

**Figure 2. Fitted response variable (Symbiodiniaceae Amplicon Sequence Variant abundance) of the GJAM model.** Tan histograms represent the distribution of the observed response variables, whereas the blue boxes show the 95% predicted response variables plotted in bins. The width of each bin stands for the x-axis values in the bin, vertical lines show the full range of predicted values within the bin, and the middle line shows the mean of the observed variables in each bin. The grey dashed line is the 1:1 line.

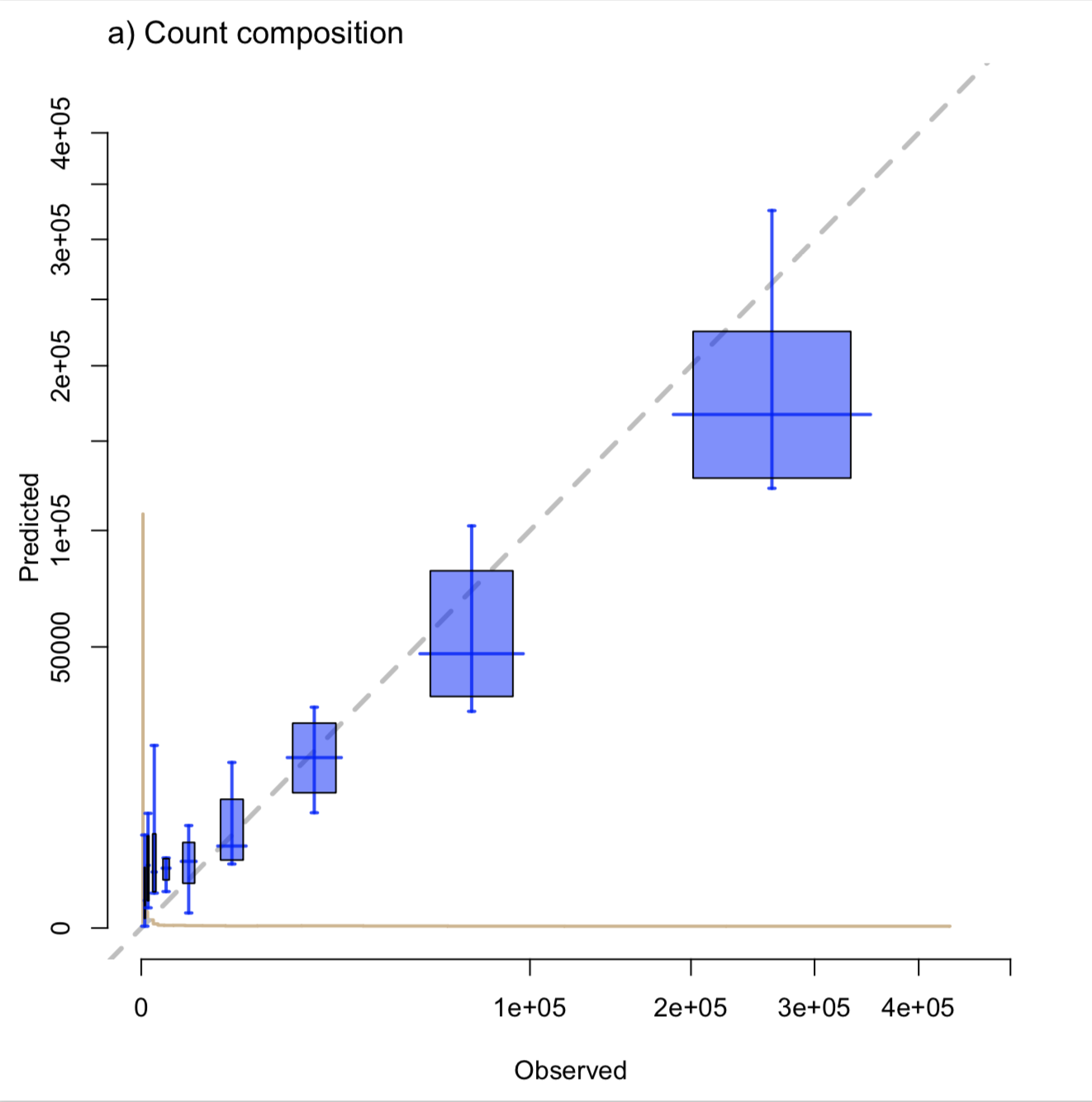

**Figure 3. Sensitivity of Symbiodiniaceae Amplicon Sequence Variants (ASVs) to *Acropora millepora* genet and experimental treatment.**

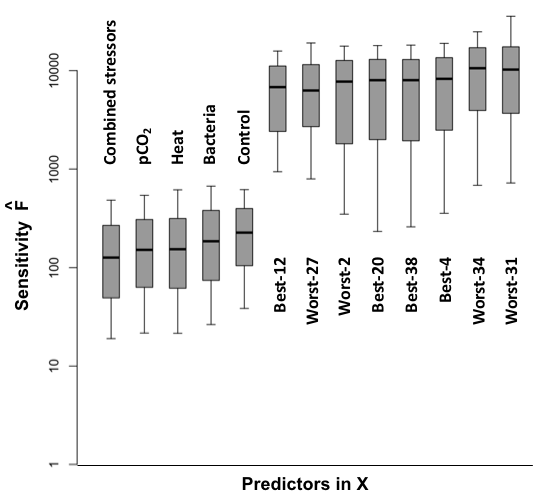

**Table 1. Pairwise Wilcoxon rank sum test of Symbiodiniaceae cell density counts by host genet.** Significant results (p < 0.05) in bold.

|  | Worst-2 | Worst-27 | Worst-31 | Worst-34 | Best-4 | Best-12 | Best-20 |
| --- | --- | --- | --- | --- | --- | --- | --- |
| Worst-27 | **0.032** | -- | -- | -- | -- | -- | -- |
| Worst-31 | 0.418 | **0.018** | **--** | -- | -- | -- | -- |
| Worst-34 | 0.052 | 0.198 | **0.033** | -- | -- | -- | -- |
| Best-4 | **0.032** | 0.829 | **0.018** | 0.152 | -- | -- | -- |
| Best-12 | 0.152 | 0.342 | **0.018** | 0.050 | 0.349 | -- | -- |
| Best-20 | 0.086 | 0.274 | **0.018** | 0.091 | 0.451 | 1.000 | -- |
| Best-38 | 0.545 | 0.171 | 0.137 | **0.032** | 0.090 | 0.437 | 0.296 |

**Table 2. Pairwise Wilcoxon rank sum test of log transformed growth data (% ∆ weight g · day^-1^) by treatment for host genets dominated by *Cladocopium* 3k and 1232 (Best-4, Best-12, Best-20, Best-38, Worst-2).** There were no significant results (p < 0.05).

|  | Combined | Bacteria | Control | Heat |
| --- | --- | --- | --- | --- |
| Bacteria | 0.89 | -- | -- | -- |
| Control | 0.62 | 0.41 | -- | -- |
| Heat | 0.37 | 0.37 | 0.21 | -- |
| *p*CO2 | 1.00 | 0.89 | 0.62 | 0.35 |

**Table 3. Total reads assigned to Symbiodiniaceae types in this study.** Total abundance of Symbiodiniaceae Internal Transcribed Spacer-2 (ITS-2) region reads per *Acropora millepora* coral fragment and # reads assigned to particular ITS-2 types.

<Link to excel file: GC Supp Table 3- totalabundancebysample biorxiv.xlsx>

**Table 4. Proportion of total reads from each sample assigned to a given Symbiodiniaceae type, the # of Symbiodiniaceae types assigned to each coral fragment, and the average # of Symbiodinianceae types identified per fragment of a given *Acropora millepora* genet.**

<Link to excel file: GC Supp Table 4- relabundancebysample biorxiv.xlsx>

**Table 5. Sensitivity of Symbiodiniaceae Amplicon Sequence Variants (ASVs) to experimental treatment and *Acropora millepora* genet on individual reefs.** Generalized Joint Attribute Models were run for genets originating from Pandora Island (Worst-27, Worst-31, and Worst-34) and Rib Reef (Best-12, Best-20, Best-38), respectively. CI = Confidence Interval; SE= Standard Error.

| **Reef** | **Factor** | **Sensitivity** | **SE** | **CI 2.5%** | **CI 97.5%** |
| --- | --- | --- | --- | --- | --- |
| Pandora | TreatmentControl | 860 | 258 | 462 | 1480 |
| Pandora | TreatmentCombined | 858 | 352 | 424 | 1740 |
| Pandora | TreatmentBacteria | 780 | 377 | 217 | 1600 |
| Pandora | TreatmentHeat | 831 | 273 | 370 | 1450 |
| Pandora | TreatmentpCO2 | 938 | 366 | 367 | 1760 |
| Pandora | GenetWorst-34 | 853 | 485 | 224 | 1950 |
| Pandora | GenetWorst-27 | 698 | 252 | 288 | 1260 |
| Pandora | GenetWorst-31 | 619 | 241 | 262 | 1180 |
| Rib | TreatmentControl | 10000 | 1780 | 6740 | 13700 |
| Rib | TreatmentCombined | 11100 | 2210 | 7350 | 15900 |
| Rib | TreatmentBacteria | 15200 | 2800 | 10400 | 21100 |
| Rib | TreatmentHeat | 12300 | 2180 | 8400 | 16900 |
| Rib | TreatmentpCO2 | 9600 | 1700 | 6640 | 13300 |
| Rib | GenetBest-38 | 14500 | 3670 | 8170 | 22000 |
| Rib | GenetBest-12 | 17600 | 2980 | 12000 | 24000 |
| Rib | GenetBest-20 | 14500 | 2360 | 10100 | 19500 |
